# Sex mosaic and a rare male in the parthenogenetic stick insect *Neohirasea japonica* (Phasmatodea, Lonchodidae)

**DOI:** 10.1101/2025.08.20.671192

**Authors:** Taisei Morishita, Tetsuo Nawa, Tomonari Nozaki

**Affiliations:** Fukue Junior High School, Tahara, Aichi 441-3615, Japan; Nawa Insect Museum, Gifu, Gifu 500-8003, Japan; Laboratory of Evolutionary Genomics, National Institute for Basic Biology (NIBB), Okazaki, Aichi 444-8585, Japan; Department of Basic Biology, School of Life Science, The Graduate University for Advanced Studies, SOKENDAI, Okazaki, Aichi 444-8585, Japan

**Keywords:** Diapheromeridae, Phasmatodea, *Neohirasea japonica*, Asexual reproduction, Gynandromorph, Sexual dimorphism

## Abstract

Parthenogenetic species that reproduce solely by females are pivotal for understanding the evolution and diversity of reproductive strategies. Rare males, often resulting from developmental errors, including chromosomal abnormalities, offer valuable insights into reproductive reversibility, although their rarity limits data on morphology, behavior, and fertility. Stick insects (Phasmatodea), with numerous parthenogenetic species, are key taxa for studying these phenomena; however, detailed analyses of rare males remain scarce. In this study, we investigated the spontaneous appearance of male and sexual mosaics within a captive colony of *Neohirasea japonica*, a widespread stick insect in Japan, where males are typically absent. In total, three individuals exhibiting male characteristics (penis and non-oviposition) were observed during the 8-years rearing. One displayed a typical male abdominal clasper and exhibited mating behavior with conspecific females, with morphological comparisons strongly suggesting that it was an *N. japonica* male. The other two individuals lacked mating behavior and were identified as sexual mosaics based on their external morphology and the presence of female reproductive systems upon dissection. This study provides foundational morphological and anatomical data on male, female, and sexually mosaic individuals in *N. japonica*. It also includes quantitative comparisons of key traits, such as the antenna-to-body length ratio, which is 0.78 in males and ranges from 0.52 to 0.53 in females. These findings establish valuable criteria for future identification of rare males and sexual mosaics in this species, ultimately contributing to our understanding of sexual trait degeneration in obligate parthenogenetic lineages.

## INTRODUCTION

Although parthenogenesis offers short-term reproductive advantages and demographic benefits, the absence of mating can lead to long-term challenges, such as the accumulation of deleterious mutations. Recent studies have revealed cryptic sexual activity in some ancient parthenogenetic lineages (Vakhrusheva et al., 2020; Boyer et al., 2021; Freitas et al., 2023), often mediated by rarely occurring males. Surprisingly, these rare males, likely arising from developmental errors, often retain sexual function (van der Kooi & Schwander, 2014; but see Nozaki et al., 2025). This persistence suggests that male-specific genes can remain functional over extended evolutionary timescales, even within asexual lineages, raising questions about the strict asexuality of long-standing parthenogenetic groups (Boyer et al., 2021).

Phasmatodea, including stick insects, is a well-studied taxon in terms of the evolution of parthenogenesis, with extensive research on the origin of parthenogenesis, geographic distributions, and degeneration of sexual traits under asexuality (Pantel, 1917; Bedford, 1978; Brock, 2000; Morgan-Richards et al., 2010; Suetsugu et al., 2023; Schwander et al., 2025). The occurrence of rare males has also been previously documented within this order (Nagashima, 2001; Morgan-Richards et al., 2010; Scali, 2013; Brock et al., 2018). The functionality of rare males is crucial for understanding the evolutionary reversibility of parthenogenesis. However, their infrequent appearance makes this research challenging. Exceptionally, in *Timema* spp., rare males in parthenogenetic species have been shown to maintain functionality (Schwander et al., 2013), and population genetic analyses of wild populations suggest the potential for occasional sexual reproduction involving these males in the past (Freitas et al., 2023). Conversely, studies on the Japanese stick insect, *Ramulus mikado*, have indicated that rare males may lose their reproductive function (Nozaki et al., 2025). Therefore, accumulating case reports with careful examinations of male functionality is essential, potentially incorporating a citizen science approach.

The genus *Neohirasea* (Phasmatodea: Lonchodidae: Necrosciinae) is widely distributed across temperate and tropical Asia (China, India, Japan, Malaysia, Taiwan, and Vietnam) with males known in most species (Table S1). In Japan, *Neohirasea japonica* has been reported as a parthenogenetic species in which only females are commonly observed (Ichikawa, 2016). To date, several instances of males with mating behavior observed in the field have been documented (Toshikiyo, 2017; Sumikawa et al., 2024, Minoshima et al., 2025); however, details regarding their reproductive functionality and the potential for sexual mosaicism, as seen in other parthenogenetic stick insects (Pijnacker, 1964), remain unknown.

In this study, we carefully examined three individuals exhibiting male-like characteristics that emerged within a captive *Neohirasea japonica* colony maintained by T. Morishita at his residence for 8 years (Fig.1, 2). Initially, we assessed mating behavior of the individuals, followed by examining their external morphology, focusing on body size, antenna length, and typical male traits, including abdominal clasper structure. We also focused on the antenna-to-body length ratio, a valuable criterion for distinguishing between the sexes within *Neohirasea*. Based on our observations, one of the three individuals was identified as male. For the remaining two, whose sex could not be definitively determined externally, their reproductive systems were assessed and compared with typical female reproductive systems.

**Fig. 1.**
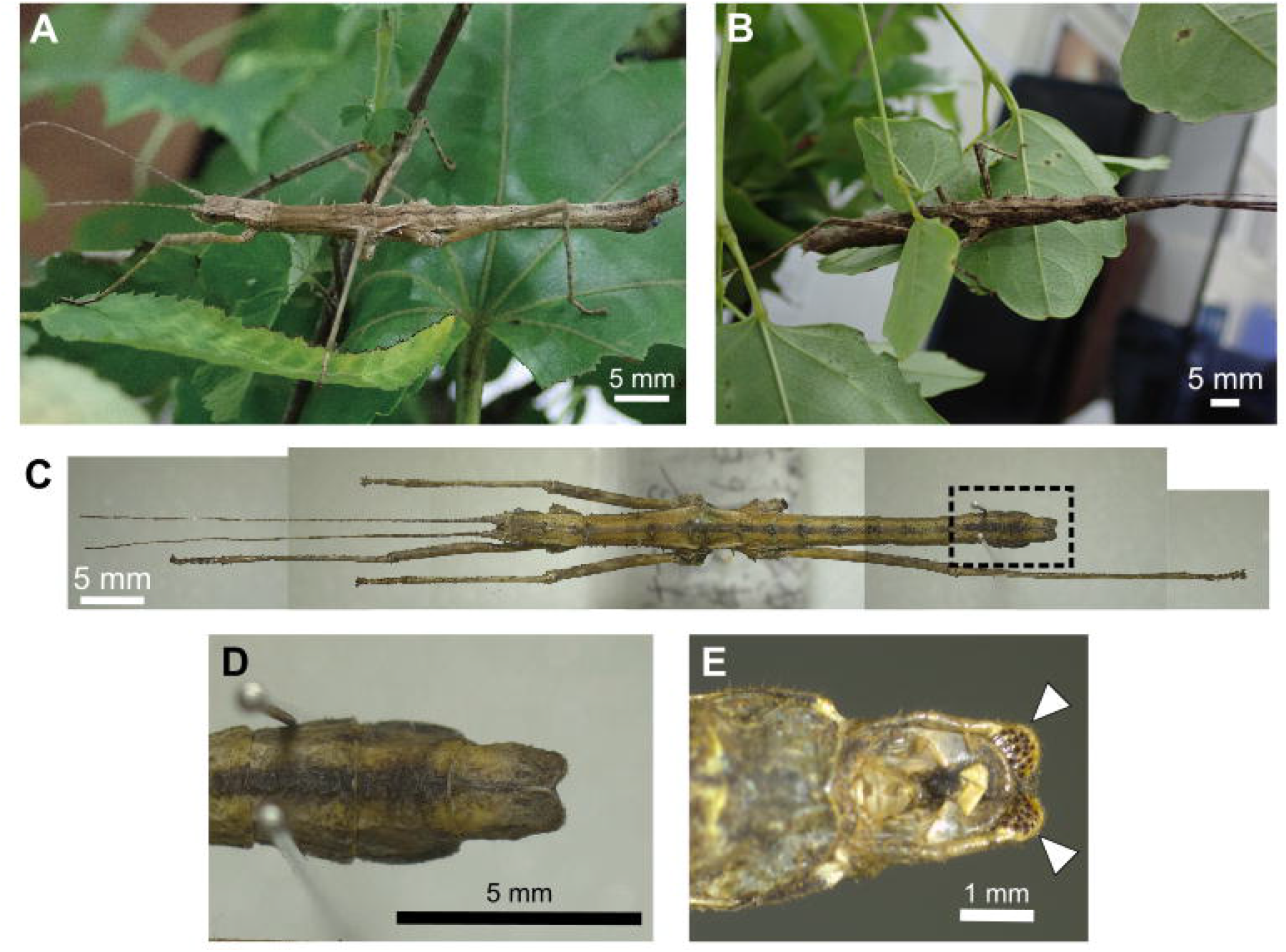
A, B. Live male and female of *Neohirasea japonica* from our captive population. (photographed by T. Nozaki at the Nawa Insect Museum). **C** Dried specimen of the male (deposited in the Nawa Insect Museum). **D, E** Enlarged views of the abdominal end (corresponding to the dashed box in **C**). **D** Dorsal view. **E** Ventral view, with the white arrowhead indicating the male-specific spines (sclerotized cuticle).

**Fig. 2.**
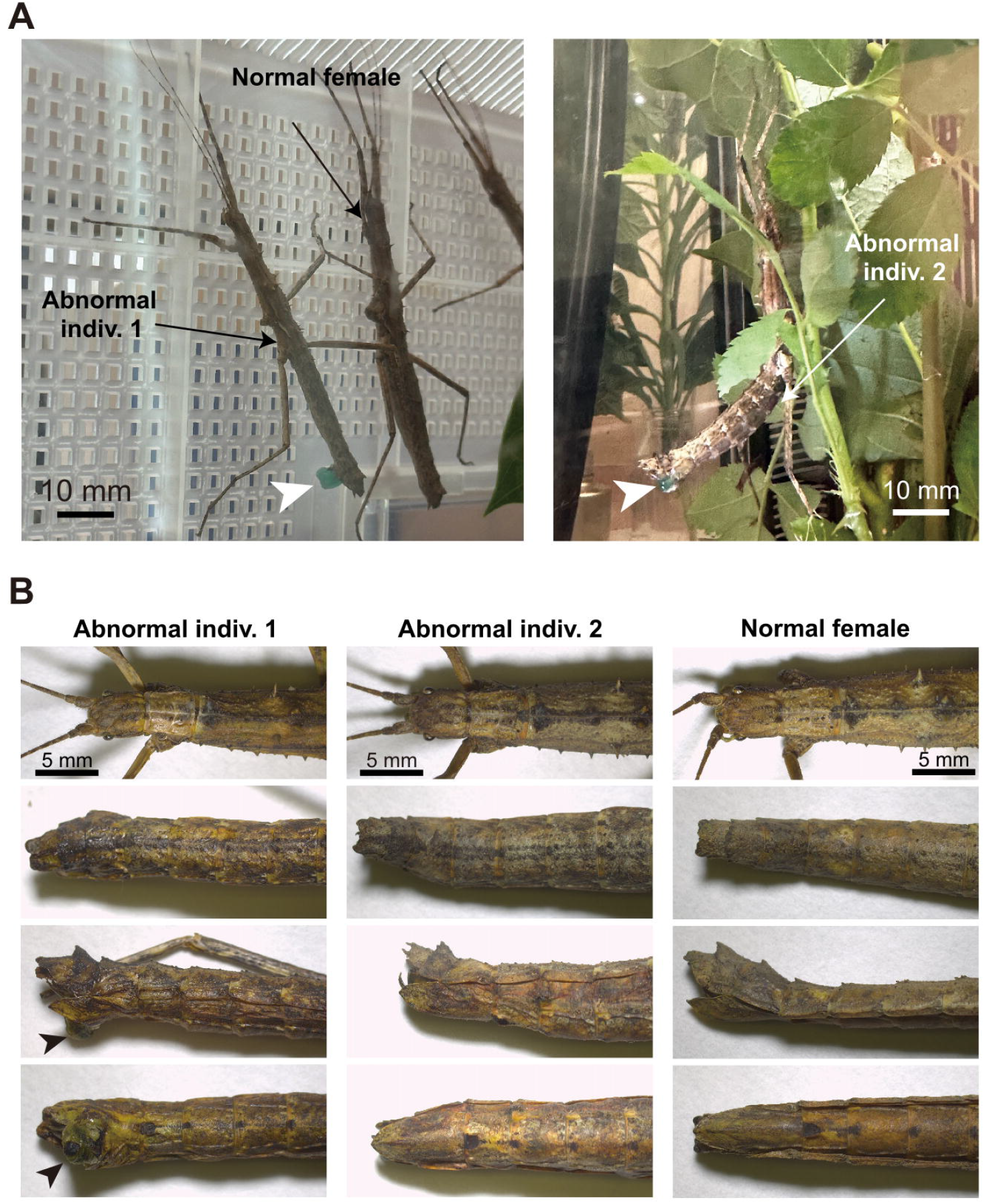
Two individuals with male-like characteristics (hereafter referred to as an abnormal individual) and a typical female from our captive population observed in 2023. Abnormal individuals possessed external genitalia (penis) but did not exhibit mating behavior. **A** Live specimen in the rearing cages. Left: Abnormal individual 1, and a typical female. Right: Abnormal individual 2 (photographed by Taisei Morishita). Arrowheads: Externally exposed penis (appearing green due to hemolymph engorgement). **B** Morphology of the head and abdominal end of abnormal individual 1, abnormal individual 2, and a typical female shown in dorsal, lateral, and ventral views. Arrowhead: exposed penis.

## MATERIALS AND METHODS

### Origin and maintenance of the captive *Neohirasea japonica* population

A captive parthenogenetic population of *Neohirasea japonica* was established from a single female collected by Taisei Morishita from a forest in Mie Prefecture in 2017. This colony has been maintained parthenogenetically for 8 years under temperature-controlled conditions (25–28 °C in summer; >10 °C in winter). Individuals were placed in plastic containers (24 cm length × 42 cm width × 29 cm height) or mesh butterfly cages (60 cm length × 60 cm width × 90 cm height) and supplied with fresh leaves of Japanese aralia (*Fatsia japonica*), Japanese spindle tree (*Euonymus japonicus*), dock (*Rumex* spp.), Japanese coral tree (*Viburnum odoratissimum*), multiflora rose (*Rosa multiflora*), chocolate vine (*Akebia quinata*), and mountain yam (*Dioscorea japonica*). Food was changed every 4–5 days. In summer, the plants were misted with tap water every 1–2 days. From summer to autumn, 300–400 eggs were collected annually. The eggs were placed on moistened floral foam (a water-absorbing sponge for fresh flowers) and stored in a sealable plastic container while maintaining the humidity. Hatching took place from January to February of the following year. The small nymphs were fed soft leaves, such as *R. multiflora* and *Alstroemeria* sp., and from the 5th instar larvae onward, they were fed the food mentioned above.

### Discovery of individuals with male characteristics

During the long-term rearing of *N. japonica*, the emergence of adult individuals exhibiting male-like characteristics (Fig. 1A, 2A; presence of a penis and absence of oviposition) was observed once in 2022 and twice in 2023. The 2022 individual underwent behavioral monitoring for several weeks before being donated to the Nawa Insect Museum. Following its display, Tetsuo Nawa prepared it as a dry specimen, which was used for subsequent morphological measurements. Individuals observed in 2023 were monitored until their natural death and then immediately frozen at −20 °C for external and internal morphological analyses.

### Morphological and anatomical observations of male-like individuals

We reviewed existing morphological data to elucidate sex differences and establish male identification criteria in *Neohirasea*. Similar to many stick insects, males in this genus possess distinctive abdominal genitalia (Ho, 2017, 2018). However, we focused on antennal length and body size, which are often reported as male traits (smaller body, proportionally longer antennae), to address potential sexual mosaics. Our review of 26 datasets across *Neohirasea* (Table S1), including data for both sexes in ten species, indicated that males consistently exhibit shorter bodies and longer antennae. Consequently, the antenna length to body length ratio (males: 0.69–1.00 [Mean ± SD: 0.82 ± 0.11]; females: 0.46–0.75 [Mean ± SD: 0.59 ± 0.09]) was considered another reliable male indicator (Table S2).

Behavioral and morphological observations were conducted to characterize male-like individuals from the captive population. We focused on the presence or absence of mating behaviors in conspecific females (Fig. 1B), expansion of the external genitalia (pennis), and egg laying. The dried specimens (the individual in 2022) were photographed using a Tg-5 digital camera (Olympus, Japan) for morphological examination. Freshly frozen specimens of the individuals in 2023 were documented using a FLOYD-4K digital camera (WRAYMER, Japan) attached to a stereomicroscope (SZ61; Olympus, Japan). Based on previous descriptions of male stick insects, we focused primarily on mating-related structures, including claspers and penises (Nagashima, 2001; Toshikiyo, 2017; Yano et al., 2021; Sumikawa et al., 2024; Nozaki et al., 2025). Additionally, antenna, body, and leg lengths were measured, and their ratios were calculated as an indicator of sex (Table S2). Abnormal individuals 1 and 2 were subsequently dissected in phosphate-buffered saline (PBS; 33 mM KH_2_PO_4_ and 33 mM Na_2_HPO_4_, pH 7.4) under a stereomicroscope using fine forceps. The presence or absence of reproductive system components, such as testes and seminal vesicles, was recorded following Matsuda (1976) and Nozaki et al. (2025). The male could not be dissected as it was preserved as a dry specimen.

To thoroughly investigate sexual characteristics, the external and internal morphologies of the females were examined. In 2024, 10 females from the captive population and 40 field-collected females, which were sampled from geographically distant locations across Honshu (Table S3), were used for detailed external observations, similar to the male-like individuals. These females lacked a visible penis-like structure and exhibited abdominal tip morphology typical of female stick insects. External morphological observations included the abdominal end and measurements of antenna, body, and leg lengths. The antennae of one field-collected female were damaged during handling and excluded from the antenna-to-body length ratio analysis. Six field-collected females from Kyoto were dissected for anatomical examination, and the presence or absence of female reproductive system components (ovaries, oviducts, spermatheca, bursa copulatrix, and accessory glands) was recorded. The remaining female individuals were either kept in captivity or returned to their collection sites.

## RESULTS

### The male-like individual observed in 2022

This individual exhibited mating attempts and penis extension, although we did not confirm penis insertion or ejaculation of seminal fluid. No further mating behavior was recorded following the transfer to the Nawa Insect Museum. Morphological examination revealed a distinct “clasper” structure at the abdominal end (Fig. 1C–E), a male-specific external organ in stick insects. Furthermore, this individual presented a smaller body length (41.0 mm) and a comparable antenna length (32.0 mm) to females (body length: 63.6 ± 3.16 mm; antenna length 33.1 ± 1.6 mm [Mean ± SD]; Fig. 3; Table S3). Its antenna-to-body length ratio was 0.78, much higher than that of typical females (0.52 ± 0.51 [Mean ± SD], Table S3). Based on its external morphology and behavior, this individual was identified as a typical male (hereafter referred to as male).

**Fig. 3.**
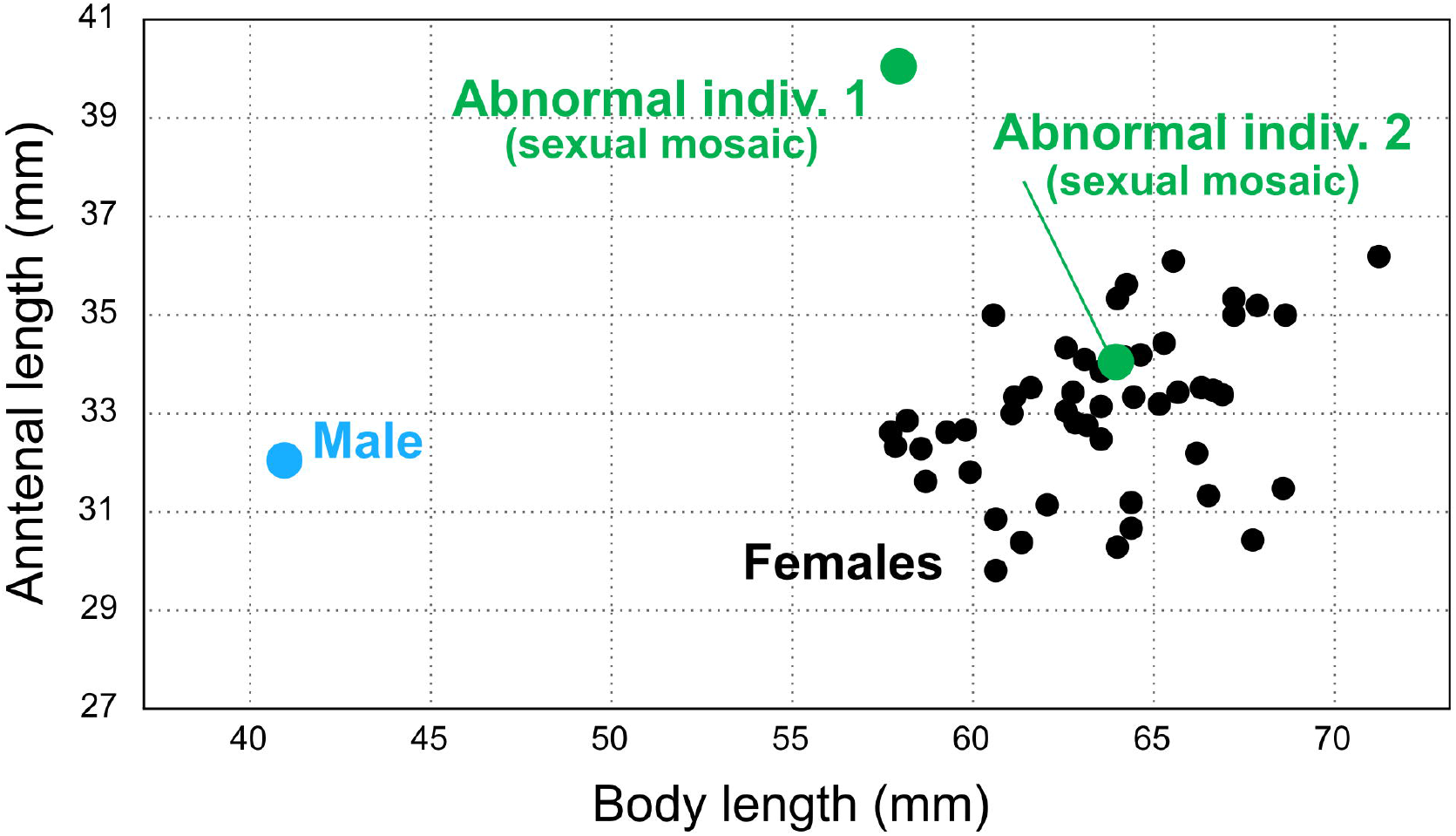
Antenna-to-body length relationship in *Neohirasea japonica* individuals used in this study. The blue dot represents the male. Green and black dots represent abnormal individuals and typical females (*n* = 49), respectively. One female out of 50 was excluded from the analysis due to damaged antennae, preventing accurate measurement.

### The two male-like individuals observed in 2023

These individuals displayed no mating behavior despite frequent penis-like structural expansion (Fig. 2A). They lacked characteristic male claspers (Fig. 2B). In contrast to typical females, which begin ovipositing within 2 weeks post-eclosion (Morishita personal observations), these individuals laid no eggs during their lifespan. These individuals were classified as sexually “abnormal” (hereafter referred to as abnormal individual 1 and abnormal individual 2). In morphological analysis, the antenna-to-body length ratios of abnormal individuals 1 and 2 were 0.69 and 0.52, respectively. These values are markedly different from the typical male ratio of 0.78, and are instead much closer to the female ratio of 0.52, with the ratio of individual 2 being identical (Table S3). Specifically, abnormal individual 1 had a slightly smaller body size (58.0 mm) but a longer antenna (40.0 mm) than typical females (Fig. 3). Individual 2 fell within the female data range for both body size (64.0 mm) and antenna length (34.0 mm) (Fig.3). We also found asymmetrical male and female features in the abdominal tip of abnormal individual 1, whereas abnormal individual 2 exhibited a consistently female-like morphology (Fig. 4A). Anatomical dissections confirmed the presence of a complete female-type reproductive system in both abnormal individuals, including a pair of ovaries, oviducts, spermathecae, bursa copulatrix, and accessory glands (Fig. 4B), along with retained degenerate oocytes.

**Fig. 4.**
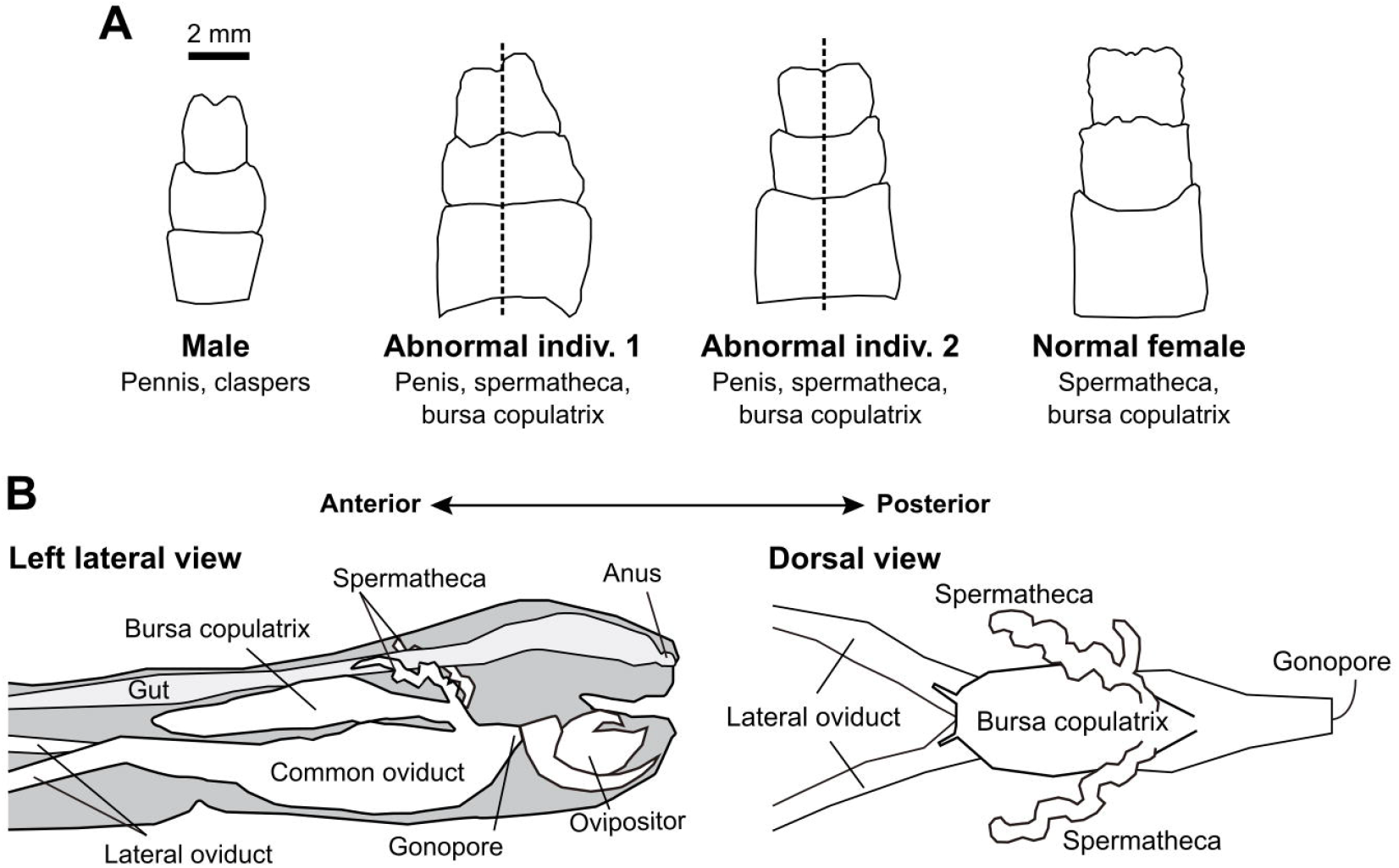
A Morphology of the abdominal end from the dorsal view, with information on the internal reproductive organs of the individuals dissected in this study. The dashed line in the diagrams indicates the midline. **B** Schematic illustration of the internal reproductive organs of a female. Abnormal individuals possessed a pair of ovaries contained oocytes, and exhibited the anatomical characteristics of females.

## DISCUSSION

In this study, we present a comprehensive analysis of three male-like individuals that emerged within a captive parthenogenetic colony of *Neohirasea japonica*. Based on our observations and the general pattern documented for this genus (Table S2), we concluded that the male-like individual collected in 2022 was a male of this species. Nevertheless, based on the behavioral, morphological, and anatomical discrepancies with a typical male, we concluded that the two individuals obtained in 2023 were likely sex mosaics (i.e., gynandromorphs) (Pereira et al., 2010). Although copulation has been observed in this species under both wild and captive conditions (Toshikiyo, 2017; Minoshima et al., 2025), future research is required to determine whether mating behavior in this species can lead to successful seminal fluid transfer from males to females as recently focused in *R. mikado* (Nozaki et al., 2025). Additionally, investigating the development of the male reproductive system and spermatogenesis would be valuable for discussing the evolutionary aspects of losing male functionality (van der Kooi & Schwander, 2014; Nozaki et al., 2025).

The long-term maintenance of our single-maternal line colony, with the emergence of only one typical male over 8 years, strongly supports the obligate parthenogenetic nature of *N. japonica* and the exceedingly rare occurrence of functional males. The controlled conditions of our study mitigated the potential for overlooking cryptic functional males, a limitation often encountered in field-based investigations. Future research should focus on elucidating the functional capabilities of these infrequent males, specifically examining the feasibility of successful mating and the paternal genetic contribution to the next generations, questions that remain largely unanswered (Schwander et al., 2013; Nozaki et al., 2025).

The occurrence of sexual mosaics reported in this study underscores developmental abnormalities in the sex determination of *N. japonica*, mirroring observations in *Carausius morosus*, where temperature-dependent incubation yields a spectrum of sexual mosaics (categorized as 0–100% males) (Pijnacker, 1964; Pijnacker & Ferwerda, 1980). This finding suggests that previous reports of rare males in *N. japonica* might have included unrecognized sexual mosaics (Toshikiyo, 2017; Sumikawa et al., 2024; Minoshima et al., 2025). Although defining males in species in which they are rarely observed presents challenges, future reports on individuals with male-like traits should include detailed descriptions of both external morphology and reproductive organ structures. Indeed, the *N. japonica* male that was identified in the present study also has the potential to be a mosaic; a male with ovarioles has been described in *R. mikado* (Nozaki et al., 2025). The accumulation of such detailed case reports will be critical for understanding the underlying developmental mechanisms leading to these abnormalities and for predicting potential disruptions in the sexual development system of the obligate parthenogenetic species *N. japonica*.

In conclusion, the present study provides the first quantitative comparative morphological data of three male-like individuals in a parthenogenetic *N. japonica* colony by identifying one male and two sexual mosaics. These findings provide crucial foundational information and valuable criteria for identifying and characterizing rare males of this species in the future, which may contribute to our understanding of sexual trait degeneration in obligate parthenogenetic species.

## ACKNOWLEDGEMENTS

We thank Nobuo Kitano, Nobuhiko Matsuo, Tatsuru Kuga, Ryo Haraki, Katsuharu Toshikiyo, Yûsuke Minoshima and Paul Brock for their helpful advice on stick insects. We also thank Tsukasa Morishita and Noriko Morishita for their support in field collection and experiments and Professor Shuji Shigenobu of NIBB for providing the experimental space and laboratory equipment. T. Morishita was financially supported by a grant from the Masason Foundation.

## SUPPORTING INFORMATION

Additional Supporting Information may be found online in the Supporting Information section at the end of the article.

**Table S1**. Morphological data of *Neohirasea spp*. from previous studies

**Table S2**. Antenna/body size ratio in *Neohirasea spp.* (from data available for both sexes, derived from Table S1)

**Table S3**. Morphological data of *Neohirasea japonica* used in this study

